# Isoform-Resolved Temporal Remodeling of Transcriptome Architecture in Monkeypox Virus

**DOI:** 10.64898/2026.10.26.761922

**Authors:** Gergely Ármin Nagy, Balázs Kakuk, Zsolt Csabai, Ákos Dörmő, István Prazsák, Daniel Růžek, Jiri Holoubek, Gábor Kemenesi, Dóra Tombácz, Zsolt Boldogkői

## Abstract

We reanalyzed previously generated replicated Oxford Nanopore direct cDNA sequencing data from six infection time points, together with cap analysis of gene expression and native direct RNA sequencing, to reconstruct full-length monkeypox virus transcripts and characterize their temporal regulation. A broad catalogue of candidate transcript models, including candidate 5′ poly(A)-associated structures, was filtered with independent 5′- and 3′-end evidence to define conservative analysis sets. Transcript-resolved profiles revealed a structured temporal program extending beyond gene-level abundance. Host-virus interaction, transcription, and nucleotide-metabolism genes contributed strongly early, whereas virion structure and morphogenesis genes dominated later. Transcription start and end site abundance profiles changed extensively over time, and many genes redistributed among alternative full-length transcript structures. Differential transcript usage was particularly prominent among host-virus interaction genes but depleted among virion structure and morphogenesis genes. Candidate 5′ poly(A)-associated transcripts also increased in relative contribution and showed locus-specific patterns. Building on our previous endpoint map, the replicated direct cDNA time course enabled temporal quantification of gene abundance, transcript-end dynamics, and isoform composition. Together, these results show that Monkeypox virus gene regulation involves coordinated remodeling of gene output, transcript boundaries, and full-length transcriptome architecture.

**Importance:** Monkeypox virus is an emerging human pathogen with the capacity for sustained human-to-human transmission, making a detailed understanding of its molecular biology increasingly important. Viral replication depends on coordinated gene expression, and defining when and how viral genes are transcribed is essential for understanding infection. Complete transcript structures provide information that gene-level measurements cannot capture because alternative RNA forms can differ in regulatory regions, coding potential, and capacity for translation. Resolving the organization and temporal use of these transcripts therefore helps reveal how the virus controls protein production and interacts with the host cell. A transcript-level view of monkeypox virus gene regulation provides a foundation for mechanistic studies of poxvirus biology and may help identify regulatory processes or viral functions relevant to antiviral intervention.

## Introduction

Poxviruses generate a highly complex transcriptional program that changes extensively over the course of infection. Gene expression is traditionally organized into early, intermediate, and late phases, with early genes predominantly supporting host interaction and establishment of infection, whereas post-replicative genes increasingly encode functions required for genome replication, transcription, virion assembly, and morphogenesis (1–3). This temporal program of Monkeypox virus (MPXV) was further characterized in subsequent studies, showing early prominence of host-interaction functions and later increases in replication, transcription, entry, and virion-associated functions (2, 4–6). This functional organization is also reflected in genome structure: the linear double-stranded DNA genome contains a conserved central region enriched for replication and structural functions, flanked by more variable terminal regions containing many host-interaction genes (7–9).

The approximately 197-kb MPXV genome contains about 200 open reading frames (ORFs) and is densely transcribed (10–12). Its transcriptional landscape includes extensive overlaps, read-throughs, alternative transcription start sites (TSSs) and transcript end sites (TESs), and polygenic RNAs, such that individual coding sequences can occur within transcripts with different boundaries and genomic contexts (13, 14). Poxviruses replicate and transcribe their genomes in the cytoplasm using largely virus-encoded machinery, and post-replicative transcription occurs within cytoplasmic viral factories (15). A characteristic consequence of this transcriptional system is RNA-polymerase slippage during initiation at many post-replicative promoters, producing non-templated 5′ poly(A) leaders (16–18), thereby generating substantial diversity in the RNA molecules associated with individual coding sequences.

Much of our current understanding of orthopoxvirus transcription derives from vaccinia virus (VACV). Early microarray and short-read studies characterized temporal gene-level expression (2, 19, 20), whereas long-read sequencing (LRS) subsequently revealed extensive alternative initiation, heterogeneous 3′ ends, polygenic RNAs, transcriptional overlap, and read-through by resolving complete RNA molecules (13, 14, 20–22). Time-resolved VACV LRS further showed that transcript architecture, as well as transcript abundance, changes during infection (22), indicating that temporal regulation extends beyond gene-level output.

Whether comparable remodeling of full-length transcript architecture occurs in MPXV remains poorly resolved. Our previous MPXV study mapped transcription start and end sites using cap analysis of gene expression sequencing (CAGE-seq) and native direct RNA sequencing (dRNA-seq) (23), revealing extensive initiation and termination heterogeneity but could not determine which 5′ and 3′ boundaries occur on the same RNA molecule or how individual transcript structures change over time. Non-templated 5′ poly(A) leaders further complicate transcript reconstruction when adapter position is used to infer native 5′ ends (16, 23, 24). Resolving these questions therefore requires long reads combined with independently supported transcript boundaries.

The aim of this study was therefore to determine how MPXV transcription changes during infection at the level of full-length transcript architecture and how these changes relate to gene-level expression, transcript-end usage, gene function, and genome organization. Using a previously generated replicated Oxford Nanopore Technologies (ONT) direct complementary DNA sequencing (dcDNA-seq) dataset spanning six stages of MPXV infection (25), we integrated long reads with independently defined 5′ and 3′ transcript boundaries to reconstruct full-length transcript models across the infection time course. We quantified temporal changes in coding-gene output, TSS and TES usage, and full-length isoform composition and examined their relationships with gene function and genomic position. We additionally reconstructed candidate transcript structures associated with 5′ poly(A) signals and followed their contribution during infection. Candidate-model discovery was separated from quantitative inference, with transcript or transcript-endpoint features retained according to analysis-specific evidence requirements. Together, these analyses extend previous endpoint mapping and gene-level temporal studies toward a transcript-resolved model of MPXV gene regulation.

## Results

### Transcript-resolved profiling and construction of the MPXV candidate transcript-model catalogue

We combined three sequencing approaches to characterize MPXV transcriptome architecture. Six infection time points: 1, 2, 4, 6, 12, and 24 hours post infection (hpi), each with three biological replicates, were profiled by ONT dcDNA-seq for transcript-model discovery and temporal quantification. CAGE-seq and dRNA-seq provided independent capped 5′-end and native-RNA 3′-end information, respectively. The standard transcript annotation workflow yielded 3,408 candidate transcript models, and a custom workflow with expanded 5′-end search settings recovered 2,266 additional structurally non-redundant models. Models unique to this set were retained as a candidate 5′ poly(A)-associated origin class.

After TES-coordinate refinement, strand-aware remapping, and deduplication, the discovery catalogue comprised 5,674 models (Fig. 1A and B; Table S1). Of these, 5,355 were assigned to viral genes and 3,067 received a main-ORF assignment. Among the 2,266 candidate 5′ poly(A)-associated models, 2,132 (94.1%) were assigned to annotated viral genes (Fig. 2B; Table S1). The genome-wide distribution of transcript models is shown in Fig. S1 and File S1.

**Figure 1.**
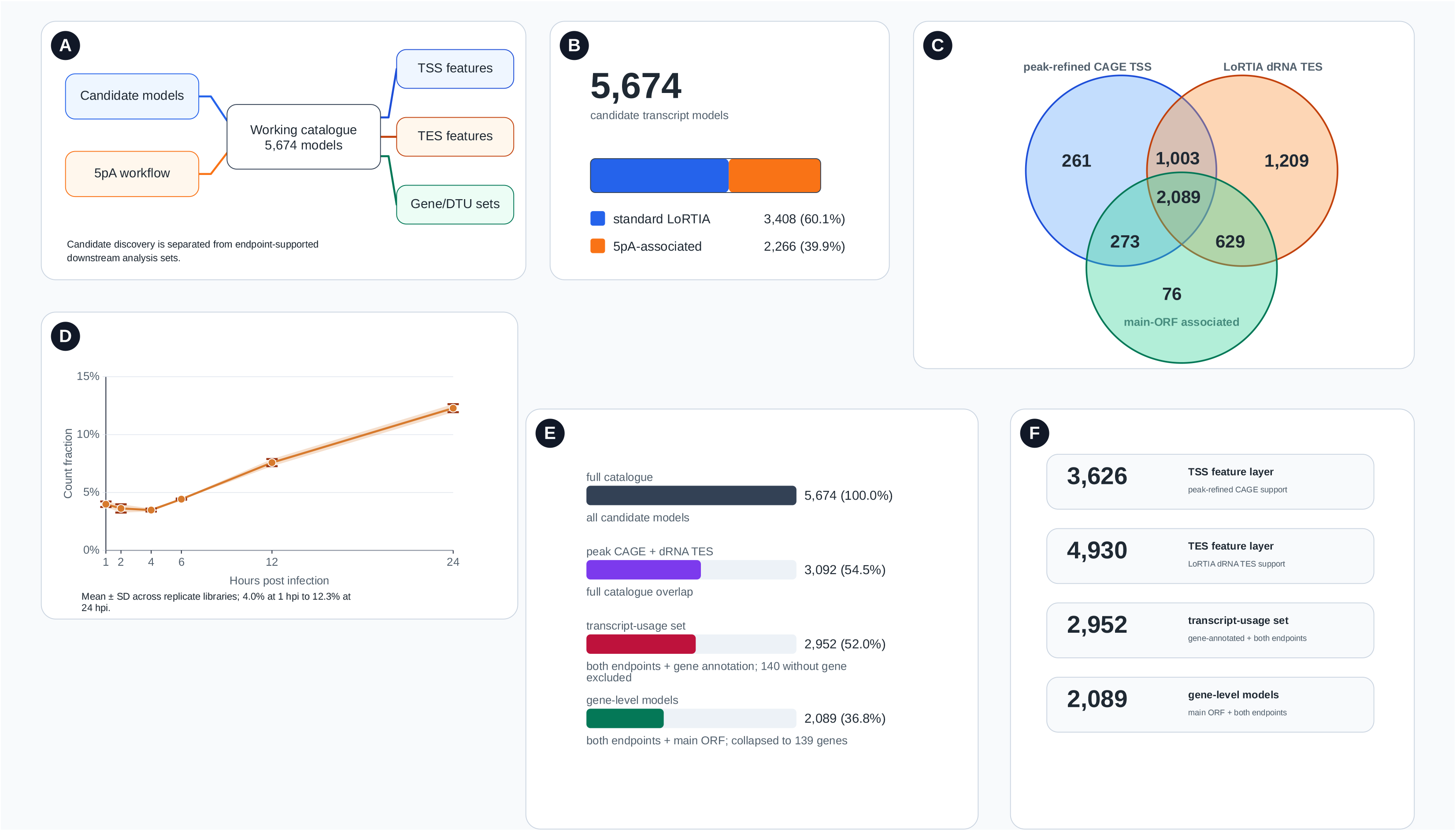
Candidate transcript-model catalogue and endpoint-supported analysis framework. A, catalogue/discovery framework. B, 5,674-model composition. C, independent endpoint and main-ORF evidence. D, candidate 5′ poly(A)-associated contribution across time. E, derivation of the 3,092-model overlap with evidence at both transcript ends, the 2,952-model transcript-usage set and the 2,089-model gene-level set. F, 3,626 TSS-qualified and 4,930 TES-qualified transcript models. Values are model counts; the endpoint layers comprise 310 TSS and 2,844 TES features (Fig. 4).

**Figure 2.**
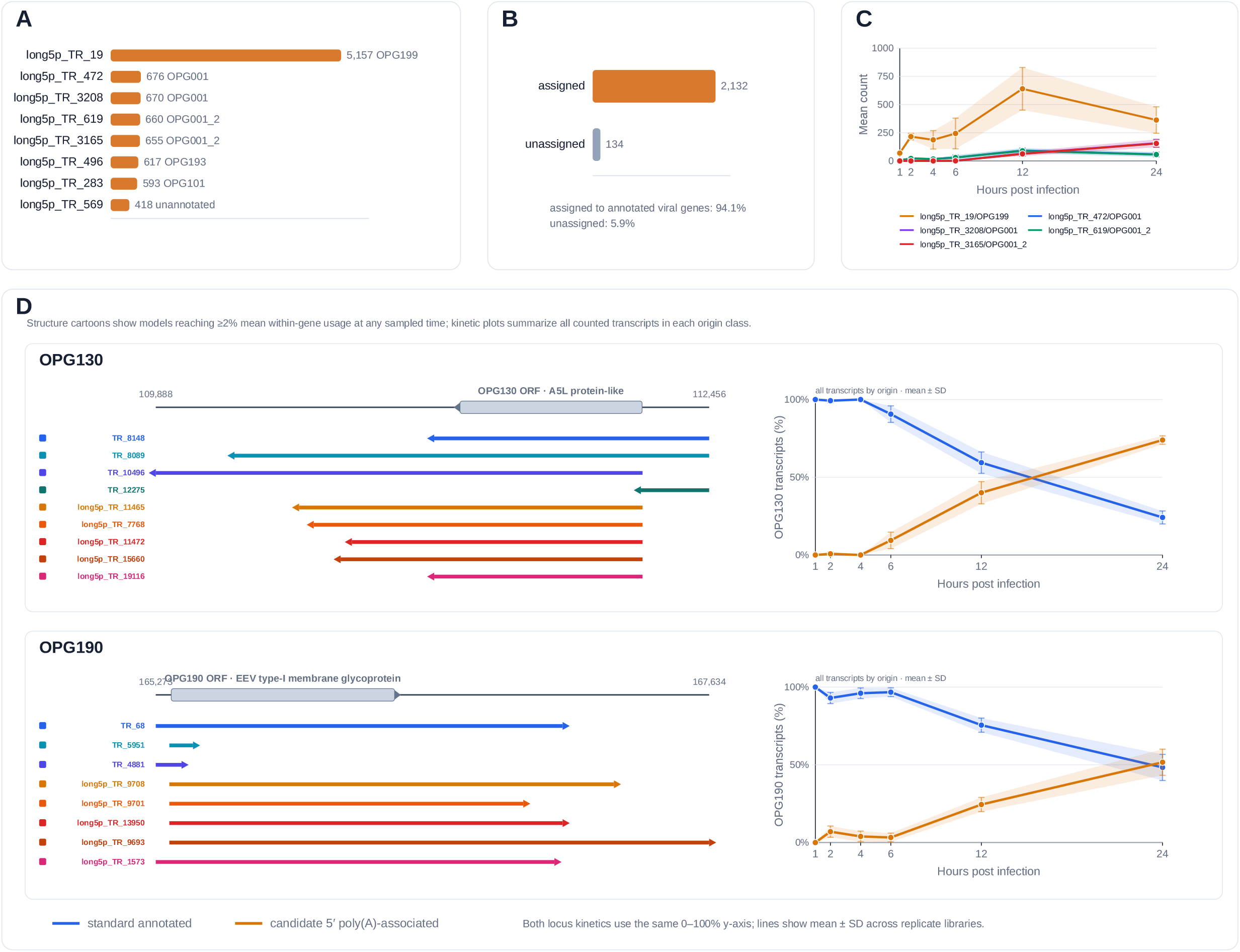
Candidate 5′ poly(A)-associated transcript models. A, Highest-count candidate 5′ poly(A)-associated models in the full transcript catalogue. Bars show total transcript counts across the 18 dcDNA-seq libraries and labels indicate assigned genes. B, Gene-assignment status among the 2,266 candidate 5′ poly(A)-associated models. C, Infection-time trajectories of the highest-count candidate models, shown as mean counts with SD across the three biological replicate libraries. D, Representative OPG130 and OPG190 loci showing temporal increases in candidate 5′ poly(A)-associated transcript usage. Transcript structures are drawn according to genomic coordinates and coloured by origin; for readability, only individual transcript models reaching ≥2% mean within-gene usage at one or more sampled time points are shown in the structural schematics. The adjacent trajectories instead include all counted transcript models assigned to each gene and show the fraction contributed by standard-annotated versus candidate 5′ poly(A)-associated models at each infection time. Lines show replicate means and error bars/bands show SD. Both loci show pronounced late increases in candidate 5′ poly(A)-associated contribution, illustrating that the transcriptome-wide increase in this origin class is concentrated at selected genes.

### Evidence-filtered layers define analysis-specific gene, endpoint, and transcript-usage sets

Independent endpoint evidence defined analysis-specific subsets of the 5,674-model catalogue. Peak-refined CAGE-seq criteria were met by 3,626 models, 4,930 had TESs corroborated by dRNA-seq, and 3,092 met both endpoint criteria (Fig. 1C, E and F). Main-ORF gene-level analysis additionally required a main-ORF assignment, retaining 2,089 models representing 139 genes and 479,869 dcDNA-seq counts (Table S2). The transcript-usage set retained 2,952 both-end-supported, gene-annotated models representing 155 genes; after restricting to genes with at least two retained models, 141 genes and 2,938 transcript features remained before DRIMSeq filtering.

Endpoint-specific analyses required only evidence for the relevant boundary. TSS analyses used adapter-correct 5′-end counts and peak-refined CAGE-supported features; TES analyses used 3′-end counts and TES features corroborated by dRNA-seq (Fig. S2A and B).

### Main-ORF gene expression undergoes widespread temporal remodeling

The evidence-filtered main-ORF gene layer revealed a structured temporal program across the infection time course (Fig. 3; Table S3). Among the 139 genes, peak-time classification assigned 29 genes to 1 hpi, 28 to 2 or 4 hpi, 36 to 6 or 12 hpi, and 46 to 24 hpi (Fig. 3A and C). A complementary classification based on the complete temporal trajectory identified 51 early-high, 39 late-rising, 17 mid-peak, 17 broad/stable, and 15 low-signal genes (Fig. 3B and D). Thus, gene expression ranged from strongly front-loaded profiles to progressive late increases, with smaller subsets showing intermediate maxima or relatively broad temporal distributions.

**Figure 3.**
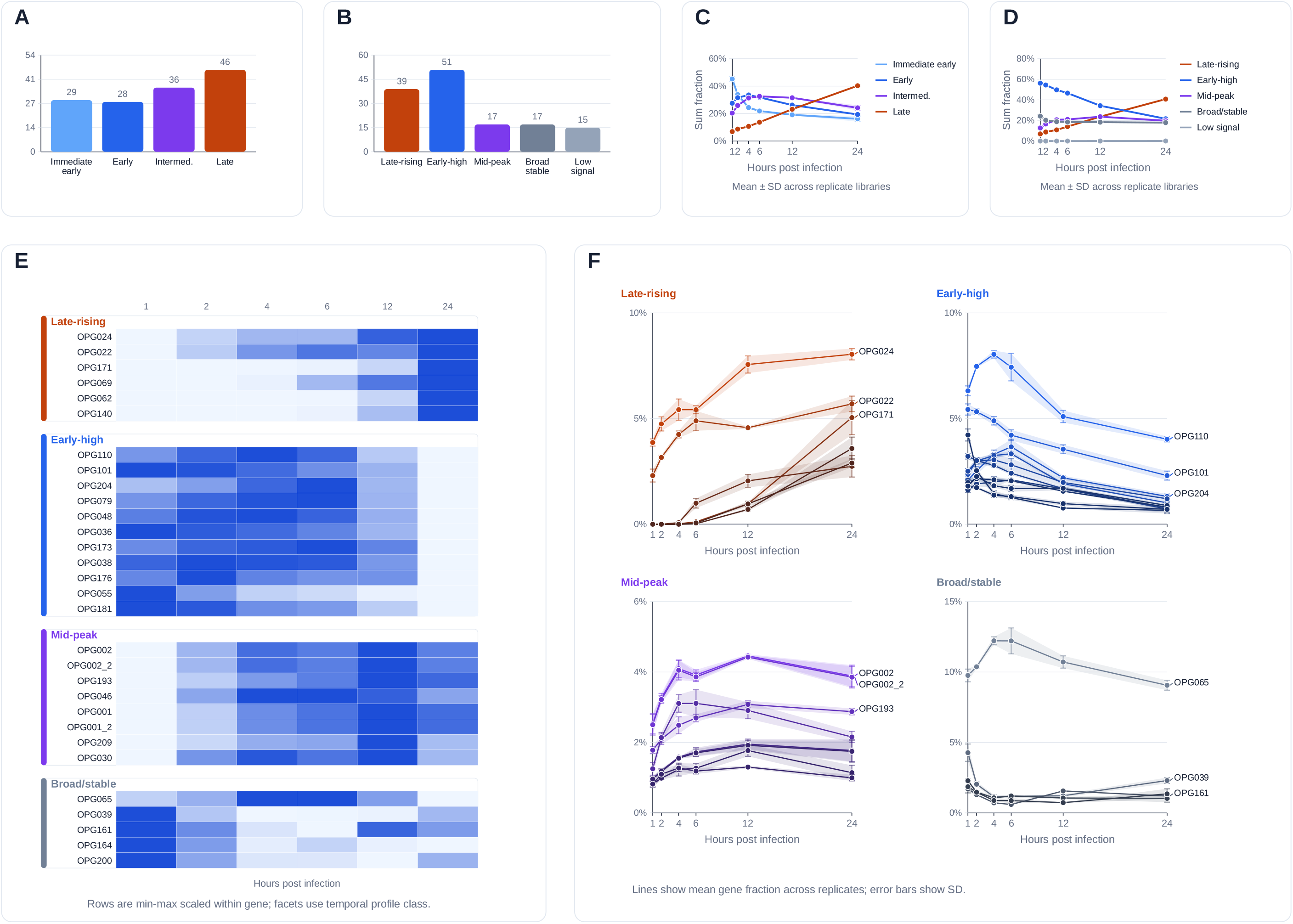
Main-ORF gene-level transcription follows a structured infection-time program. A, operational peak-time bins: 29 genes with a 1-h peak, 28 with a 2–4-h peak, 36 with a 6–12-h peak and 46 with a 24-h peak. These are observed sampling-time maxima rather than classical kinetic-gene categories. B, profile classes: 39 late-rising, 51 early-high, 17 mid-peak, 17 broad/stable and 15 low-signal. C, mean relative-abundance trajectories of the peak-time bins. D, mean relative-abundance trajectories of the profile classes. E, row-scaled heatmap of representative genes from each profile class. F, mean gene fractions over infection time for representative genes, grouped by profile class; shading shows SD across the three replicate libraries.

Highly represented orthopoxvirus genes (OPGs) occurred in contrasting temporal classes. OPG024 and OPG022 combined high cumulative transcript abundance with late-rising profiles, whereas OPG110 and OPG101 were both highly represented and early-high (Figs. S3 and S4). Other representative late-rising genes included OPG171, OPG069, OPG062, OPG140, OPG077, and OPG138, while OPG204, OPG079, OPG048, OPG036, OPG173, and OPG038 showed early-high trajectories. Highly represented genes were therefore distributed across early-high and late-rising profiles, rather than forming a single abundance-dominant temporal group. Representative gene-level heatmaps and trajectories illustrate these contrasting temporal programs (Fig. 3E and F).

Formal differential-abundance testing supported this widespread temporal regulation. After requiring at least five counts in at least three libraries, 123 of the 139 main-ORF genes were retained for DESeq2 likelihood-ratio testing. This analysis used infection time as a six-level factor and tested each gene’s full temporal profile against a no-time-effect model, rather than asking only whether a specific pair of time points differed. Of the retained genes, 108 (87.8%) showed a significant infection-time effect (Table S4; Fig. S5A and B). Temporal regulation therefore involved most sufficiently expressed main-ORF genes rather than a small subset.

Comparison with published MPXV gene-level timing profiles showed broad agreement for the late structural program and for several early genes, but some individual loci differed. OPG024 was classified as early by Picarone et al. (5) but was late-rising in the present analysis, whereas OPG071 was classified as intermediate/late by Picarone et al. but was early-high here (Supplementary Results S10, Fig. S6, Table S5). These differences may reflect the different experimental systems, including VK2/E6E7 cells sampled at 2, 24, and 48 hpi in Picarone et al. compared with CV-1 cells sampled at 1, 2, 4, 6, 12, and 24 hpi in the present dataset.

### Supported TSS and TES features reveal extensive endpoint diversity and temporal remodeling

The 5′ endpoint-specific layer comprised 310 peak-refined CAGE-supported TSS features representing 455,530 dcDNA-seq counts and mapping to 160 genes (Fig. 4A; Table S6). Eighty-eight genes (55.0%) had at least two supported TSS features (Fig. 4B and C). OPG040 illustrates this complexity with five supported TSS features showing distinct temporal abundance profiles (Fig. 4D).

**Figure 4.**
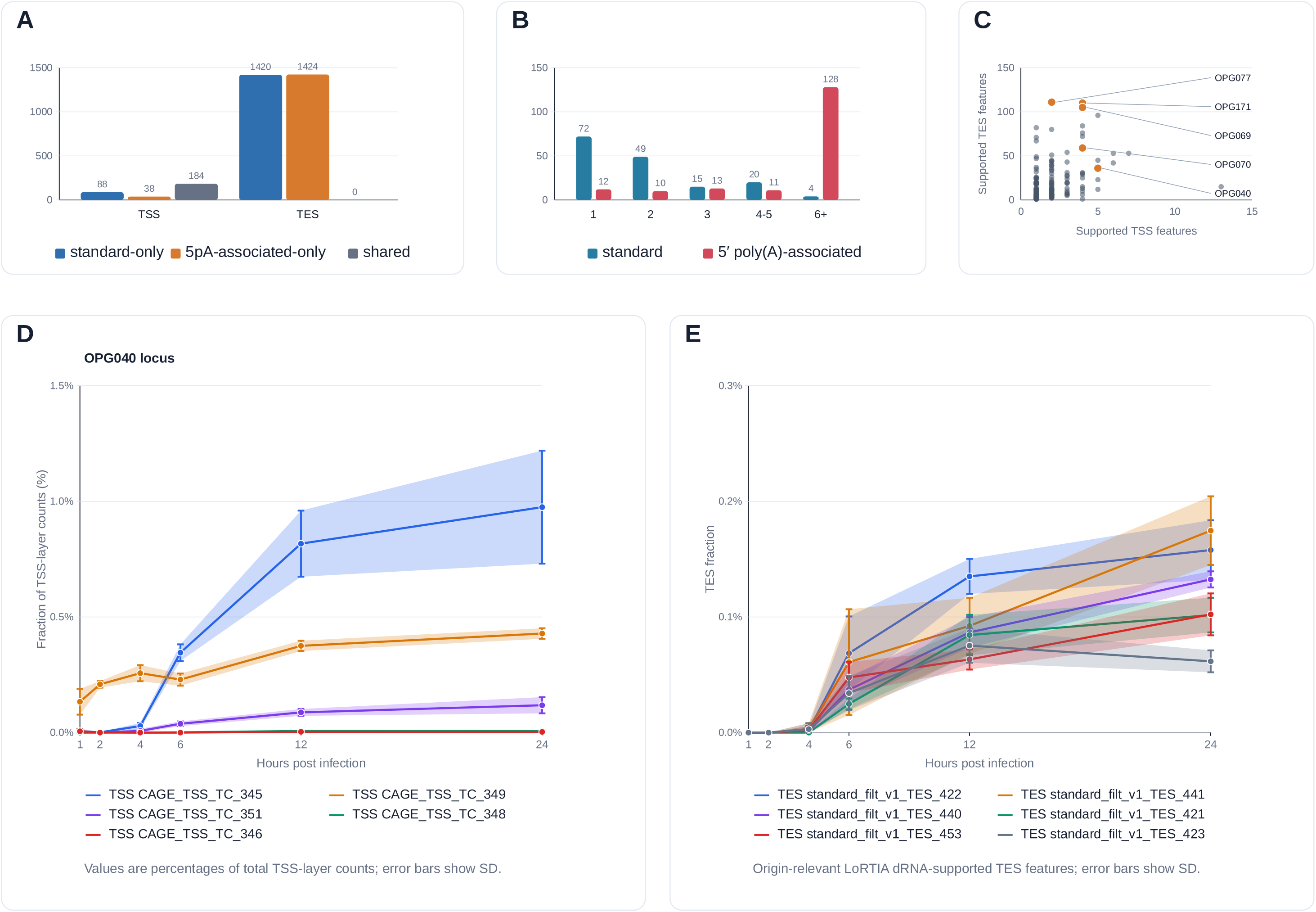
Supported TSS and TES features reveal extensive MPXV endpoint architecture. A, 310 peak-refined CAGE-supported TSS features and 2,844 origin-relevant LoRTIA dRNA-supported TES features. B–C, endpoint multiplicity/diversity. D, OPG040-associated TSS trajectories as percentages of total TSS-layer counts. E, OPG069–OPG071-region TES trajectories as percentages of total TES-layer counts. Panels D and E show endpoint-layer abundance, not within-gene endpoint-usage proportions.

The 3′ endpoint-specific layer comprised 2,844 origin-relevant TES features corroborated by dRNA-seq, representing 538,027 dcDNA-seq counts and mapping to 174 genes (Fig. 4A; Table S7). Of these genes, 162 (93.1%) had at least two supported TES features, indicating extensive 3′-end heterogeneity (Fig. 4B and C). The overlapping OPG069–OPG071 region provides an example of local TES complexity (Fig. 4E; Fig. S7).

Endpoint-feature abundance was extensively remodeled during infection. After count filtering, DESeq2 likelihood-ratio testing identified significant time effects in 178 of 221 tested TSS features (80.5%) and 693 of 985 tested TES features (70.4%; Table S4). Fig. 4D and E show fractions of total endpoint-layer signal rather than within-gene endpoint usage.

Genome-wide strand-resolved maps illustrate the temporal redistribution of supported 5′-end signals at 4 and 12 hpi (Fig. 5A and B) and the corresponding 3′-end signals at the same time points (Fig. 6A and B); six-time-point views are provided in Figs. S8 and S9.

**Figure 5.**
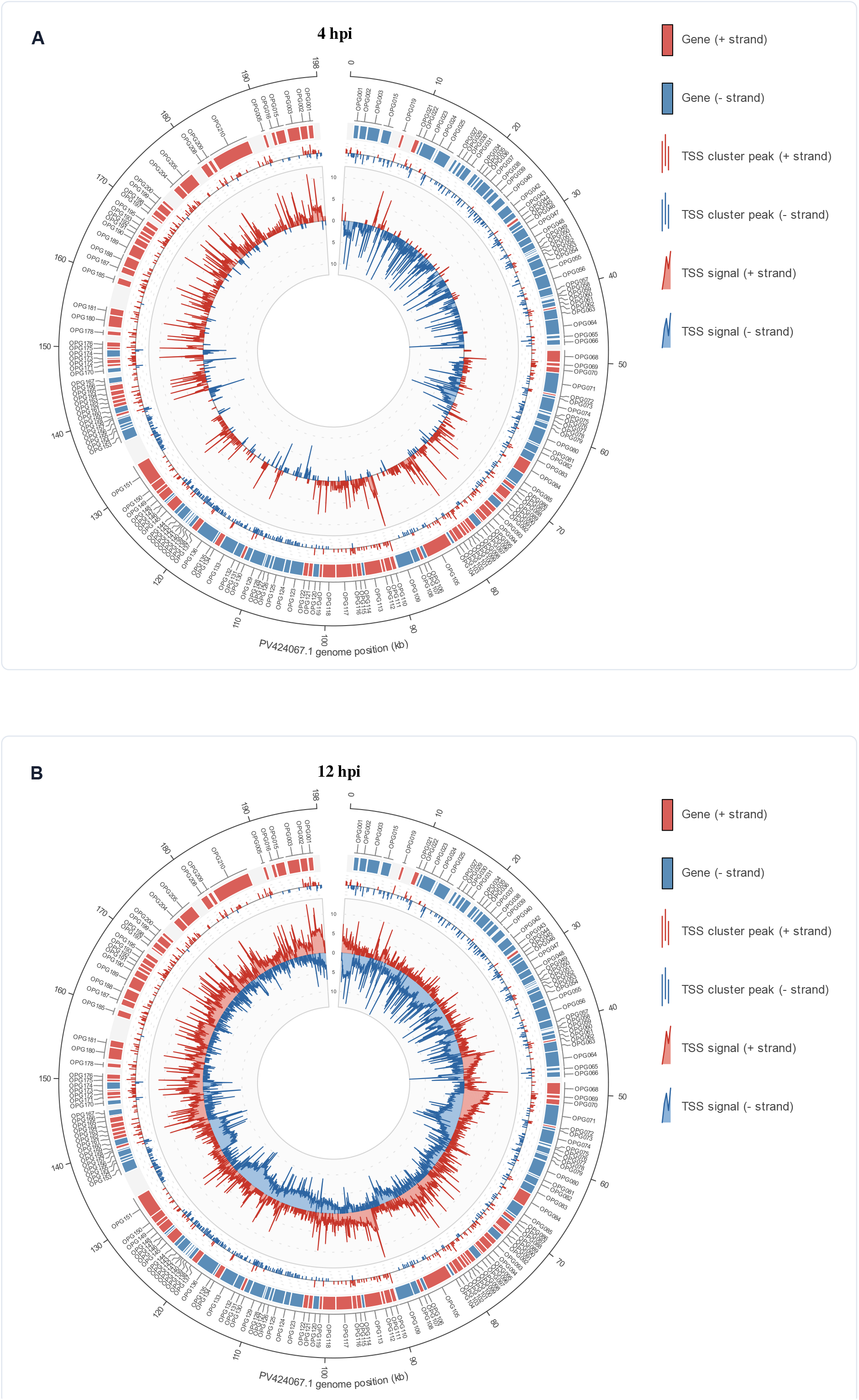
Genome-wide distribution of MPXV 5′-end signal at selected infection times. Panels A and B show 4 and 12 hpi with coding features, peak-refined CAGE-supported TSS peaks and adapter-correct dcDNA-seq 5′-end signal displayed as log2(signal + 1).

**Figure 6.**
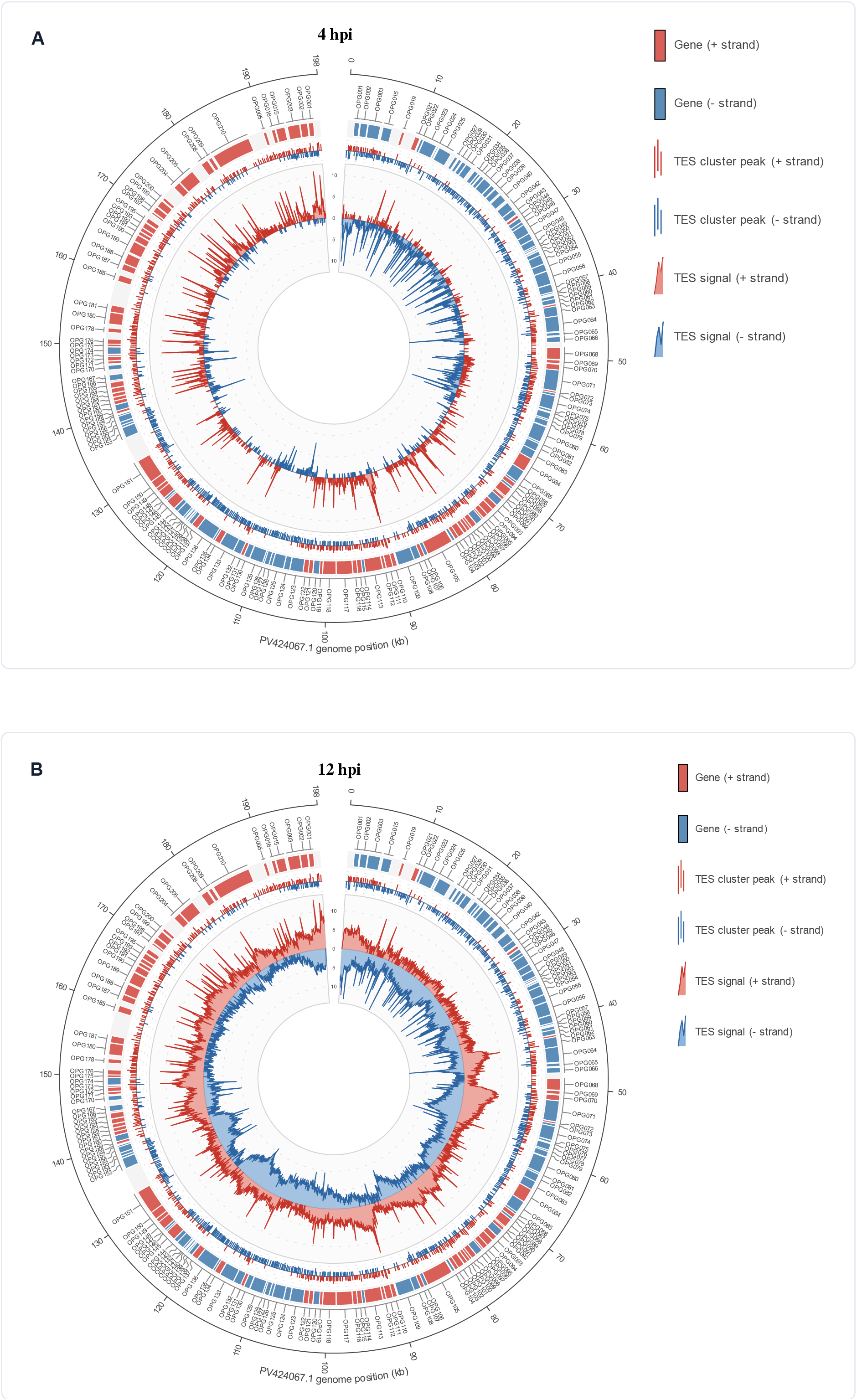
Genome-wide distribution of MPXV 3′-end signal at selected infection times. Panels A and B show 4 and 12 hpi with coding features, origin-relevant LoRTIA dRNA-supported TES peaks and adapter-correct dcDNA-seq 3′-end signal displayed as log2(signal + 1).

### Candidate 5′ poly(A)-associated transcripts show gene-specific late enrichment

These models accounted for 39.9% of the structural complexity of the discovery catalogue but contributed a substantially smaller fraction of dcDNA-seq transcript abundance. Their mean contribution to transcript counts increased from 4.0% at 1 hpi to 12.3% at 24 hpi, accompanied by a reciprocal decrease in standard-workflow models from 96.0% to 87.7% (Fig. 1D; Fig. S10).

This increase was not distributed uniformly across the transcriptome. Gene-level analysis revealed pronounced temporal shifts at a subset of loci (Fig. S11). For example, the corresponding fraction of OPG187 increased from 3.3% at 1 hpi to 31.9% at 24 hpi, while OPG054 showed a particularly strong late increase between 12 and 24 hpi. Such changes were considerably larger than the decreases observed at most genes, indicating that the transcriptome-wide late enrichment was driven disproportionately by selected loci rather than by a uniform shift across viral genes.

Their abundance was also concentrated among selected individual transcript models (Fig. S11). The highest-count model, long5p_TR_19, was assigned to OPG199 and contributed 5,157 counts across the 18 dcDNA-seq libraries (Fig. 2A–C). At the gene level, however, the temporal contribution of candidate 5′ poly(A)-associated transcripts varied markedly among loci. OPG130 and OPG190 illustrate two pronounced late shifts: both were dominated by standard-annotated transcripts early in infection, whereas the candidate 5′ poly(A)-associated fraction increased substantially at later time points, becoming the predominant origin class for OPG130 and approaching an approximately equal contribution for OPG190 by 24 hpi (Fig. 2D). These locus-level changes reinforce that the overall temporal increase in candidate 5′ poly(A)-associated transcripts was driven disproportionately by selected genes rather than by a uniform transcriptome-wide shift.

### Differential transcript usage reveals time-dependent isoform redistribution

Differential transcript usage (DTU) was used to test time-dependent redistribution among both-end-supported transcript models within genes. Starting from 2,952 gene-annotated, both-end-supported models, DRIMSeq retained 85 genes and 493 transcript features after abundance and proportion filtering. Gene-level testing identified 40 genes with BH-adjusted P values below 0.05, while stageR confirmed 93 transcript features representing 39 genes at an overall false-discovery rate of 5%. Requiring both gene-level DRIMSeq significance and at least one stageR-confirmed transcript yielded 37 confirmed DTU genes (43.5% of the 85 modeled genes; Fig. 7A–C; Table S8A and B). Of the 93 stageR-confirmed features, 90 belonged to these 37 genes; the remaining three were associated with OPG019 and OPG170, which did not meet the gene-level BH-adjusted significance criterion.

**Figure 7.**
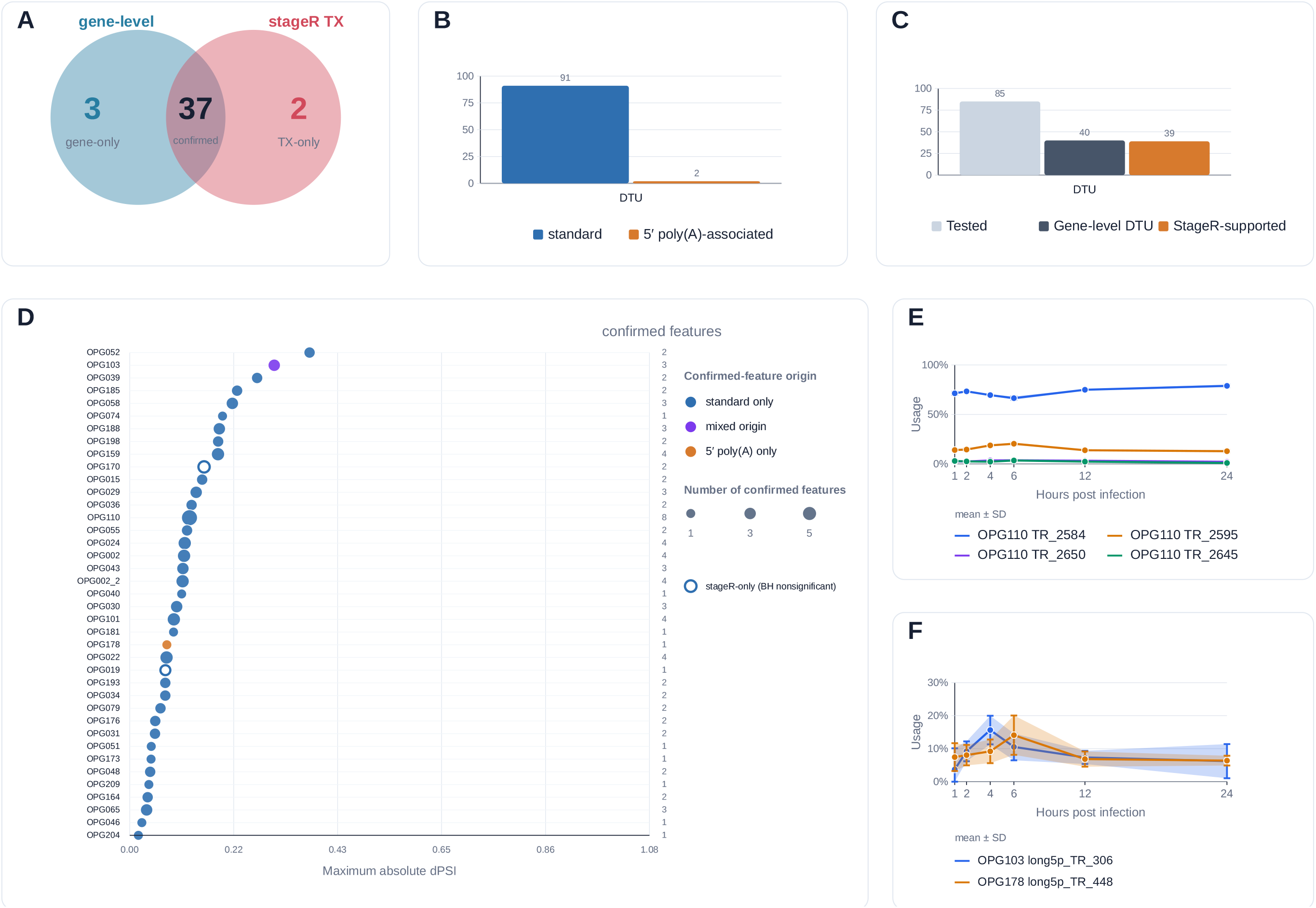
A, Overlap between genes significant by DRIMSeq after BH adjustment and genes carrying at least one stageR-confirmed transcript feature (three DRIMSeq-only, 37 shared, and two stageR-only genes). B, Origin of the 93 stageR-confirmed transcript features (91 standard and two candidate 5′ poly(A)-associated models). C, Numbers of genes tested by DRIMSeq (85), significant at the gene level (40), and carrying stageR-confirmed transcript features (39). D, All 39 genes carrying stageR-confirmed transcript features, ranked by maximum absolute dPSI. Point size indicates the number of stageR-confirmed transcript features per gene, and point colour indicates their origin (standard-only, candidate 5′ poly(A)-associated-only, or mixed). Filled points represent the 37 genes meeting both criteria; hollow points identify OPG019 and OPG170, which carried stageR-confirmed features but did not meet the independent gene-level DRIMSeq BH-adjusted significance criterion. E–F, Representative within-gene transcript-usage trajectories across infection time; error bars and shading indicate SD across the three biological replicates.

DTU occurred across distinct temporal expression programs. OPG110 showed highly significant redistribution among multiple transcript models (gene-level P = 1.07 × 10^-54), with eight stageR-confirmed features, whereas the late-rising OPG024 locus also showed strong DTU (P = 1.38 × 10^-56) with four confirmed features. OPG039 combined strong statistical support with a larger proportional shift: TR_8009 changed by dPSI = -0.265 between 1 and 12 h (gene-level P = 2.12 × 10^-148). The distribution of maximum absolute dPSI values and representative OPG110 isoform trajectories are shown in Fig. 7D and E.

The clearest large-amplitude redistribution occurred at OPG052. TR_7734 decreased from approximately 65.9% of modeled transcript usage at 1 h to 29.1% at 24 h, while TR_7617 increased from 0% to approximately 49.1%. Of these, TR_7734 and TR_7731 were stageR-confirmed; TR_7734 had a maximum absolute dPSI of 0.374. The overall OPG052 DTU signal was highly significant (gene-level P = 1.77 × 10^-33), indicating marked late redistribution of transcript usage within the locus.

Candidate 5′ poly(A)-associated models also contributed to confirmed DTU (Fig. 7F). Pairwise dPSI summaries identified the time contrasts with the largest proportional changes (Fig. S13), with several strong shifts involving comparisons with 24 h, consistent with substantial late-stage remodeling of transcript composition.

### Functional organization links timing with transcript architecture

We assigned all 179 reference OPG features to broad NCLDV-compatible functional categories through manual curation, integrating comparative sequence similarity, conserved domain annotations, and published functional evidence (Table S9). Of these, 169 assignments agreed with Picarone et al. (5), while the remaining ten differed and are detailed in Table S10. Within the evidence-filtered main-ORF set of 139 genes, the largest classes were host-virus interactions (50 genes), virion structure and morphogenesis (34), and transcription and RNA processing (22), with the remainder distributed among uncharacterized, DNA replication/recombination/repair, nucleotide-metabolism, miscellaneous, and other metabolic functions (Fig. 8A; Table S11A).

**Figure 8.**
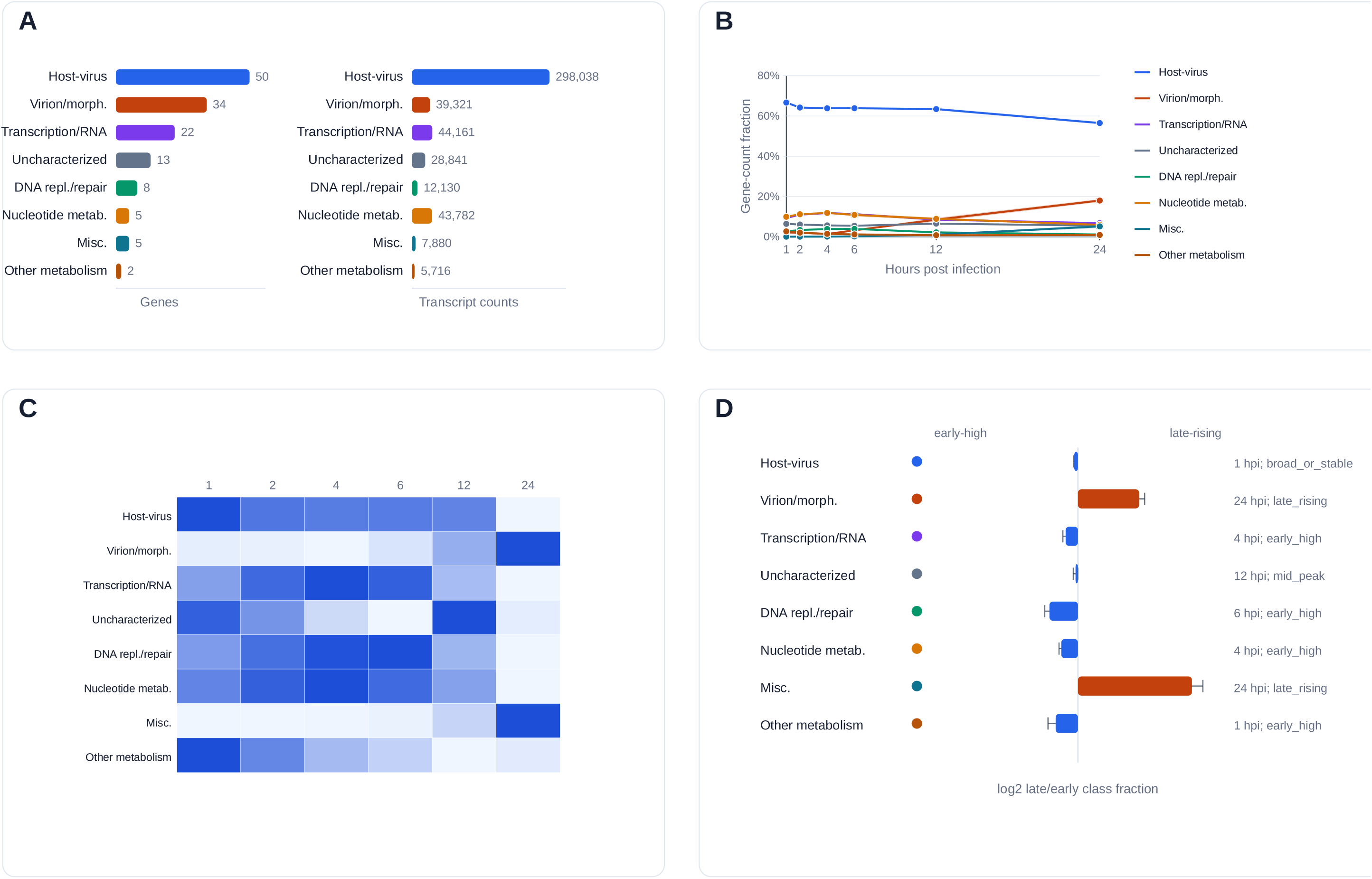
Functional classes define distinct endpoint-supported MPXV transcriptional programs. A, gene numbers and transcript-count signal for 139 main-ORF genes. B, functional-class trajectories as fractions of main-ORF gene-layer counts. C, row-scaled trajectories. D, class-level timing with publication-formatted profile labels.

Functional classes showed distinct temporal profiles in their relative abundance trajectories and class-level heatmaps (Fig. 8B and C). Virion structure and morphogenesis genes were predominantly late: the class peaked at 24 hpi and 20 of 34 genes were late-rising. Transcription/RNA-processing and nucleotide-metabolism classes peaked earlier, whereas DNA replication/recombination/repair showed an intermediate peak but an early-high overall profile (Fig. 8D). Comparison with VACV-WR showed high coding-feature similarity across the central region (Fig. 9A). This pattern of greater central conservation was consistent across all seven orthopoxvirus comparisons (Supplementary Results S9, Fig. S14). Virion structure/morphogenesis and transcription/RNA-processing genes were concentrated predominantly in the central region, whereas host-virus interaction genes were more frequent toward the variable termini (Fig. 9B).

**Figure 9.**
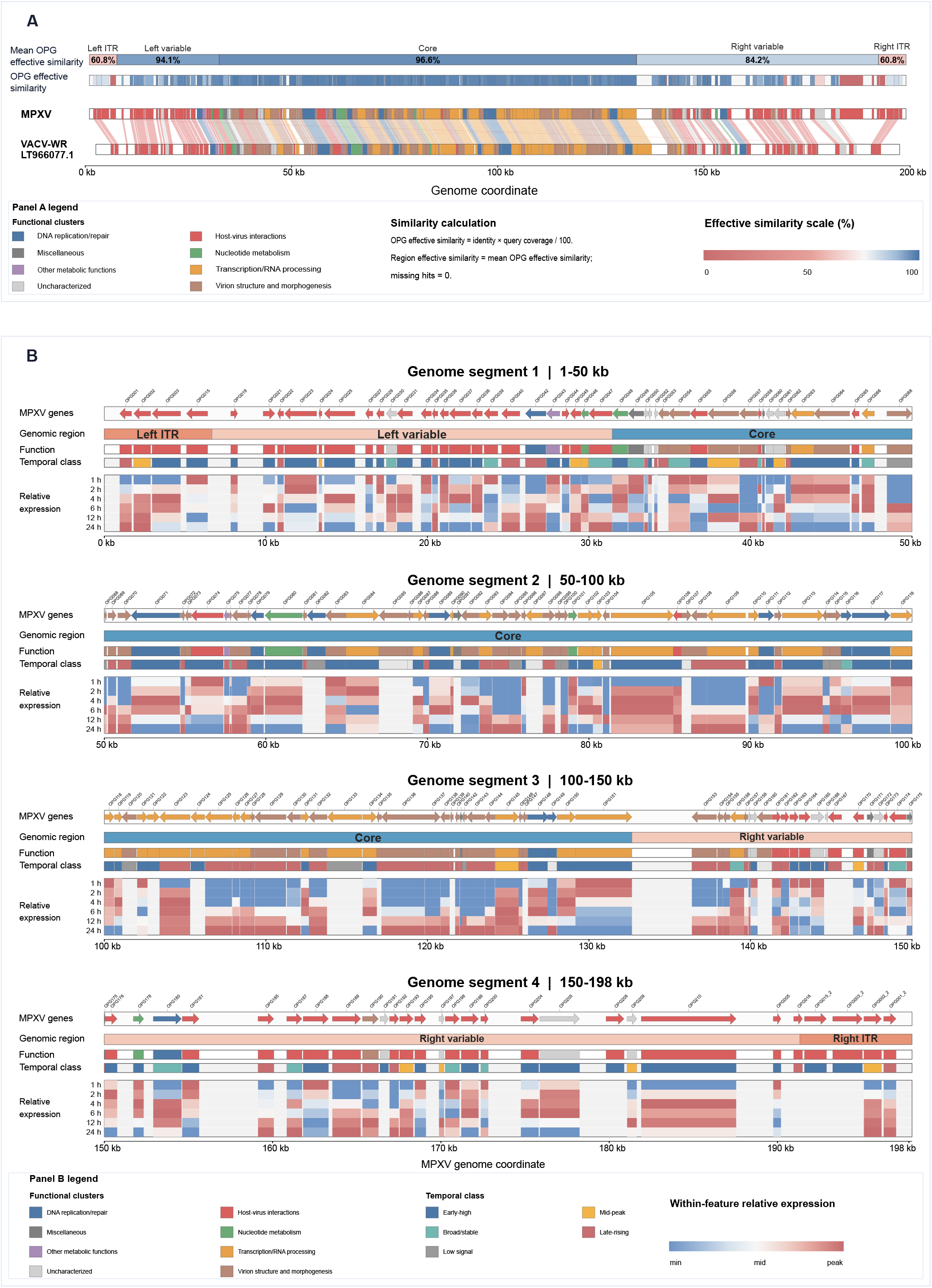
Coding-feature conservation and temporal transcriptional architecture of MPXV relative to VACV-WR. A, genome-wide comparison of MPXV with VACV-WR showing mean OPG effective similarity per genomic region, OPG-level effective similarity, and the aligned coding features of both genomes. B, the MPXV genome in four segments, each showing coding features, genomic region, functional class, temporal class and row-scaled relative expression at 1, 2, 4, 6, 12 and 24 hpi. OPG effective similarity = alignment identity × query coverage/100; regional values are mean OPG effective similarity with missing hits assigned 0.

Candidate 5′ poly(A)-associated contribution, TSS and TES multiplicity, terminal localization, temporal profile, and confirmed DTU varied among functional classes (Fig. 10A through E). To summarize these patterns, we performed a functional-association analysis using feature-specific eligible gene sets, with omnibus tests followed by class-specific one-versus-rest Fisher exact tests (Fig. 10F; Table S11B). Functional class was associated with temporal profile and confirmed DTU, whereas high TSS and TES diversity did not show significant class-level associations under the selected thresholds. In the post hoc tests, late-rising profiles were enriched among virion structure and morphogenesis genes and depleted among host-virus interaction genes, while early-high profiles were depleted among virion structure and morphogenesis genes. Confirmed DTU was enriched among host-virus interaction genes and depleted among virion structure and morphogenesis genes when restricted to DRIMSeq/stageR-testable genes. These associations indicate functional differences in transcript-composition remodeling but should be interpreted cautiously because of unequal class sizes, sparse contingency-table cells, and duplicated or copy-related loci.

**Figure 10.**
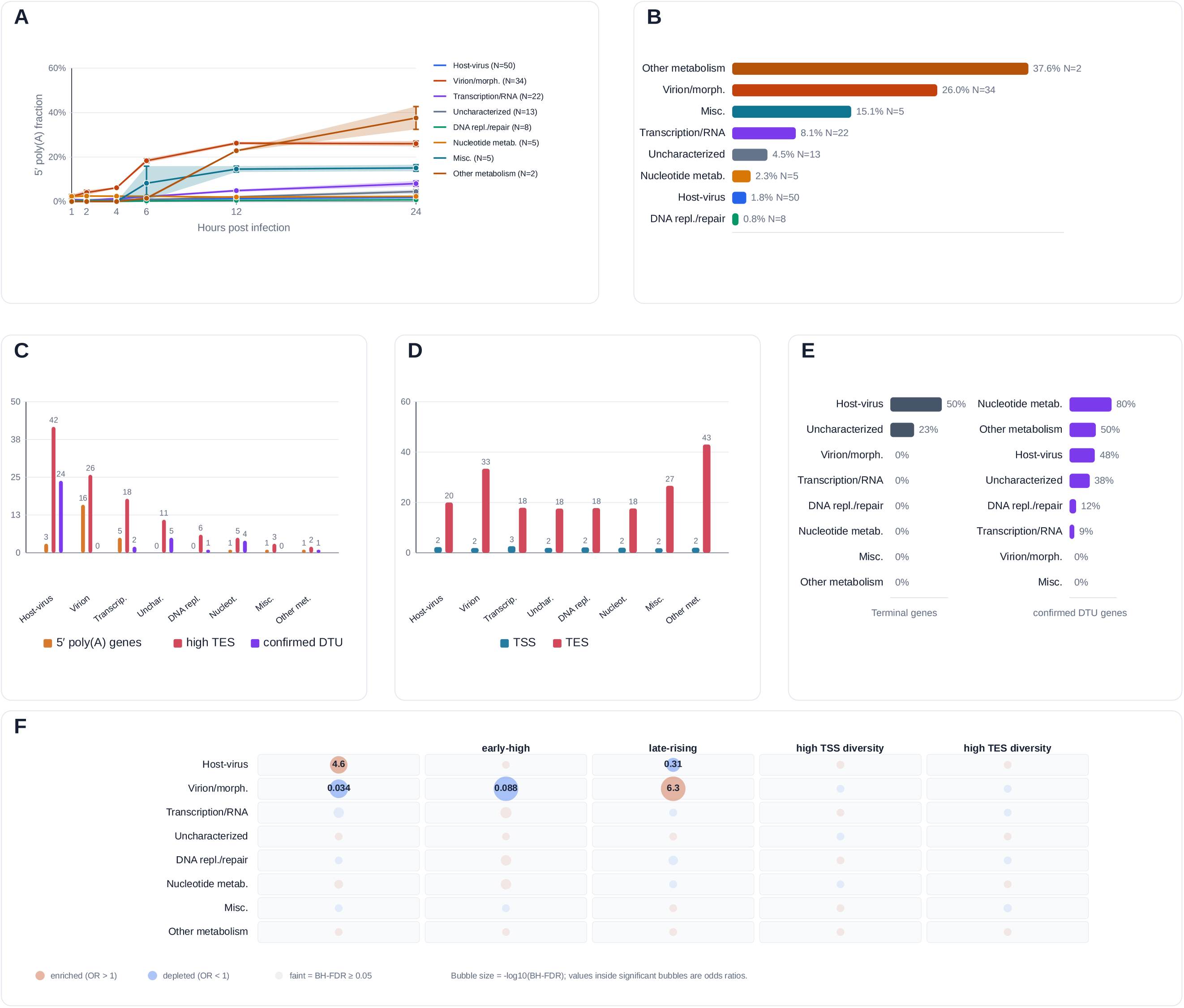
Functional classes differ in endpoint architecture and transcript usage. A, candidate 5′ poly(A)-associated contribution. B, classes ranked by the 24-hpi values from panel A. C, high TSS diversity = ≥4 supported TSS features and high TES diversity = ≥6. D, mean supported TSS/TES feature counts. E, terminal localization, defined as within 25 kb of either genome end, and confirmed-DTU fractions. F, Fisher exact association matrix.

## Discussion

This study provides a transcript-resolved view of the temporal MPXV transcriptional program by combining a replicated six-time-point dcDNA-seq series with independent CAGE-seq and dRNA-seq endpoint evidence. Analysis-specific evidence filters separated coding-gene output, transcript-end dynamics, and transcript composition among models supported at both ends. The principal advance over previous gene-level MPXV studies is the ability to follow individual transcript models across infection. To our knowledge, this is the first full-length transcript-model-resolved temporal analysis of MPXV infection.

The temporal organization of MPXV gene expression broadly agreed with previous orthopoxvirus studies, including recent MPXV transcriptomic analyses (5, 6), especially for the late structural program. Some gene-level differences were observed, such as OPG024 and OPG071, and may reflect differences in cell type, sampling times, sequencing strategy, and transcript quantification in densely transcribed regions. The denser early sampling in the present CV-1 dataset allowed a more detailed characterization of early-high and intermediate temporal profiles. These patterns should nevertheless be interpreted as relative expression trajectories rather than discrete promoter-defined kinetic classes, as illustrated by VACV studies showing that transcription from some early promoters can be reactivated after DNA replication (26).

Transcript architecture changed alongside gene abundance. Alternative 5′ and especially 3′ boundaries were widespread, and most tested TSS and TES features showed significant temporal effects. These findings extend our previous MPXV endpoint map, which identified initiation and termination complexity but could not connect both ends on individual molecules or quantify their temporal trajectories (23). Similar complexity has been described in VACV, including alternative and within-ORF TSSs, polycistronic RNAs, read-through, and extensive transcriptional overlap (13, 14, 22, 27). The extensive 3′ heterogeneity is also consistent with orthopoxvirus termination biology, in which post-replicative transcripts are particularly heterogeneous and overlapping (14).

DTU provides direct evidence that temporal regulation extends beyond changes in total gene output. Nearly half of genes with sufficient multi-isoform expression for modeling showed confirmed time-dependent redistribution among both-end-supported models, including both standard and candidate 5′ poly(A)-associated transcript structures. These changes reflect shifts in the relative abundance of RNA isoforms carrying different combinations of 5′ and 3′ boundaries rather than changes in coding-locus abundance alone. For example, OPG052 showed a marked switch in the dominant modeled transcript structure between early and late infection, illustrating how the prevalent architecture of a locus can change during infection. More broadly, DTU reveals a layer of temporal regulation not captured by gene-level abundance. However, DTU does not by itself identify whether a change in transcript-model usage is driven by altered initiation, altered termination, transcript stability, or a combination of these processes.

Candidate 5′ poly(A)-associated models formed a distinctive component of this remodeling. Non-templated 5′ poly(A) leaders arise by RNA-polymerase slippage during poxvirus transcription initiation (16,18), and their position between the detected adapter and genome-aligned sequence can cause genuine leader-containing molecules to fail standard adapter-position criteria. We therefore treated models recovered only by the expanded 5′ workflow as a candidate origin class rather than as individually proven leader-containing RNAs. Their contribution increased over infection, extending the poly(A)-associated endpoint signals previously observed in MPXV (23).

The temporal increase in these models is consistent with production of 5′ poly(A) leaders during post-replicative VACV transcription (14, 28). Such leaders can enhance translation in infected cells (28), but their presence is not definitive evidence of late transcription or efficient translation. Poly(A) leaders have also been reported on specific early transcripts, and long VACV poly(A) leaders are frequently uncapped (29, 30).

Gene-level analysis further showed that this increase was not uniform across the transcriptome but was driven disproportionately by a subset of loci showing pronounced, often late-onset increases in candidate 5′ poly(A)-associated transcript fractions, including OPG187, OPG130, and OPG190 (Fig. 2D; Fig. S10). This heterogeneity is consistent with a combination of general post-replicative transcriptional processes and gene-specific regulation rather than a uniform property of all late transcripts. Because our structural classification cannot distinguish genuine alternative initiation from transcriptional slippage or other sources of 5′ heterogeneity, and because cap status and translation were not measured here, the mechanistic and functional consequences of these transcript structures require direct validation.

Main-ORF assignment allowed transcript-level abundance to be summarized at the protein-coding gene level but reflects predicted coding potential rather than direct evidence of protein production. For this assignment, we linked each transcript to the most upstream reference-sense ORF that it fully covered, providing an operational coding-locus assignment for structurally polycistronic and read-through poxvirus RNAs (13, 22). However, assigning transcripts to a common main ORF does not necessarily imply identical translational potential. VACV ribosome profiling has identified translation from annotated starts, downstream methionines, short ORFs, and unannotated initiation sites (27). Moreover, transcripts assigned to the same main ORF may differ in 5′ leader length and structure, upstream AUG or uORF content, and initiation context, potentially influencing translation (27, 31). In yeast and human cells, alternative 5′-untranslated region (UTR) isoforms have been shown to alter translational output independently of total mRNA abundance (32, 33), and VACV studies likewise support a regulatory role for viral 5′ architecture (27,28). Alternative MPXV 5′ structures may therefore influence translation efficiency or initiation-site choice, a possibility that can now be tested directly.

Functional annotation showed that transcript-resolved remodeling was organized by viral gene function and genome position rather than being uniformly distributed across the MPXV genome. Virion structure and morphogenesis genes formed the clearest late functional program, with a 24-h class-level peak and enrichment of late-rising profiles. These genes, together with transcription/RNA-processing and other conserved replicative functions, were concentrated outside the terminal 25-kb regions. In contrast, host-virus interaction genes were the only major functional class substantially represented near the genome termini and showed the strongest confirmed-DTU enrichment among genes eligible for transcript-usage modeling. Thus, the analysis links three layers of MPXV organization: conserved central coding content, late structural gene output, and transcript-composition remodeling in the more variable host-interaction compartment. These class-level associations remain functional summaries rather than mechanistic proof, because class sizes were unequal, some contingency tables were sparse, and duplicated or copy-related loci may not be fully independent.

LRS resolves transcript architecture directly, but technical artifacts can inflate apparent diversity. cDNA workflows may generate false ends, truncated molecules, or chimeric structures, whereas dRNA-seq remains limited by incomplete molecule recovery and reduced 5′-end precision (24, 35, 36). We therefore combined adapter-aware detection and replicate/read-support criteria with independent CAGE-seq and dRNA-seq endpoint evidence, restricting transcript-level and DTU analyses to models supported at both ends. These measures reduce, but cannot eliminate, uncertainty in individual transcript models.

Several limitations remain. CAGE-seq and dRNA-seq were generated from mixed-time-point material and therefore define endpoint locations but do not independently quantify temporal dynamics. The high-MOI CV-1 system may not reproduce temporal behavior in primary tissues, other cell types, lower-dose infection, or in vivo. Transcript abundances were normalized within the viral transcriptome, thus the analysis does not establish absolute RNA abundance per infected cell. Transcript-to-ORF assignment likewise indicates coding potential rather than protein production.

Overall, MPXV transcription is organized at the levels of coding-locus output, transcript-end dynamics, and within-gene transcript composition. Replicated long-read temporal quantification with independent endpoint evidence revealed extensive remodeling across all three layers and prioritizes candidate loci for testing how transcriptome architecture influences translation, protein production, and virus-host interactions.

## Materials and Methods

### Source datasets and sequencing procedures

This study integrates previously generated MPXV transcriptomic datasets; no new wet-lab experiments were performed. Experimental procedures are described in Kakuk et al. (25) for dcDNA-seq and Nagy et al. (23) for CAGE-seq and dRNA-seq. Briefly, CV-1 cells were infected with MPXV isolate MPXV_NRL 4279/2022 at a multiplicity of infection of 5. RNA was collected at 1, 2, 4, 6, 12, and 24 hpi in three biological replicates for dcDNA-seq. A pooled mixed-time-point RNA sample was used for dRNA-seq, and CAGE-seq libraries came from a complementary endpoint-mapping dataset. Work with infectious material was performed under BSL-4 conditions at the National Laboratory of Virology, University of Pécs.

### Bioinformatic processing and catalogue construction

Statistical analyses were performed in R 4.4.3 using data.table (37), GenomicRanges (38), rtracklayer (39), ggplot2 (40), edgeR (41), DESeq2 (42), DRIMSeq (43), stageR (44), and supporting packages.

Previously generated Nanopore alignments to MPXV reference genome PV424067.1 were reused. CAGE-seq reads were aligned to the same reference with STAR v2.7.11b (45), using a maximum intron length of 1,000 nt and gene-count output.

Stranded Nanopore alignments were processed with LoRTIA v0.1 (46). TSS, TES, and intron features were merged within 10 nt and retained when detected in at least two samples with at least five total supporting reads, then assembled into candidate models with Transcript Annotator. Standard LoRTIA 5′-end parameters were check_in_soft=15, check_in_match=10, and check_from_alignment=3; the additional workflow with expanded 5′-end search settings used 100, 15, and 100, respectively.

For 5′-end quality control in the expanded-search workflow, the strand-appropriate soft clip plus 10 nt of mapped sequence was inspected for adapter sequence and a non-templated A/T tract. Reads were accepted when the adapter-to-boundary distance was <4 nt; at 4 nt the intervening sequence had to be entirely strand-appropriate A/T, and at >4 nt A/T had to cover at least 75% of the interval. Reads with accepted 5′ support or a correct strand-appropriate 3′ LoRTIA adapter tag were retained for transfrag construction. This QC contributed to the expanded-search transfrag set but did not itself define the identity of models classified as unique to this workflow. Implementation details are provided in the deposited scripts.

CAGE-derived TSS peaks and dRNA-derived TES peaks were identified with TSSr (47). dRNA TES clusters were canonicalized by reference and strand. Candidate transcript 3′ coordinates within 10 nt of an origin-relevant TES peak were reassigned to that peak before catalogue remapping and deduplication.

For 3′ endpoint support, models were evaluated against their corresponding standard or expanded-search LoRTIA run. TESs were considered dRNA-supported when dRNA evidence was present in the feature metadata or counts, with candidate coordinates matched on the same reference and strand within 10 nt of the supported cluster edge.

The standard and expanded-search catalogues were merged using strand-aware structural similarity, allowing ≤15-nt differences at TSSs/TESs and ≤2 nt at splice junctions. Gene labels were transferred to compatible unassigned expanded-search models. After TES refinement and deduplication, the catalogue contained 5,674 models; candidate 5′ poly(A)-associated models were defined by their origin in the expanded-search workflow and do not imply per-model confirmation of a non-templated leader.

### Transcript quantification, coding-locus assignment, and analysis-specific filtering

For temporal quantification, dcDNA-seq transfrags were assigned to catalogue models within 10 nt at the TSS, 15 nt at the TES, and 2 nt at each splice junction.

Each model was assigned to the most upstream fully covered reference-sense ORF; models missing no more than 5% of ORF length at the 5′ edge remained eligible.

All downstream analyses used analysis-specific endpoint criteria. Models were considered gene-annotated when assigned a gene label and main-ORF-associated when they received a positive main-ORF assignment.

Peak-refined CAGE support retained interval-supported models when the matched CAGE cluster was ≤25 nt wide; for wider clusters, the transcript TSS also had to lie within 15 nt of the dominant peak. This retained 3,626 models for TSS-supported transcript sets.

The TES layer used origin-relevant LoRTIA TES features corroborated by dRNA-seq. Both-end transcript analyses retained only models whose 5′ and 3′ boundaries met the corresponding CAGE-seq and dRNA-seq criteria.

Main-ORF analysis retained both-end-supported models with a main-ORF assignment and summed their counts by ORF, yielding 2,089 models across 139 genes. DTU analysis retained both-end-supported, gene-annotated models; after excluding single-model genes, 141 genes and 2,938 transcript features remained.

TSS analyses used adapter-correct dcDNA-seq 5′-end counts aggregated by peak-refined CAGE clusters. TES analyses used corresponding 3′-end counts aggregated by origin-specific LoRTIA TES features corroborated by dRNA-seq. All temporal inference used the 18 dcDNA-seq libraries from six time points and three biological replicates.

### Differential abundance and differential transcript usage

Raw integer count matrices were constructed separately for main-ORF genes, TSS features, and dRNA-corroborated LoRTIA TES features. Before differential-abundance analysis, each layer required at least five counts in at least three of the 18 libraries.

Temporal differential abundance was tested with DESeq2 using infection time as a six-level factor. Likelihood-ratio tests compared the full time model with an intercept-only model; Benjamini–Hochberg-adjusted P≤0.05 defined significance (48). Pairwise Wald contrasts were exported as supporting results.

Within-gene transcript usage was tested with DRIMSeq across the 18 libraries. Abundance and proportion filtering reduced the dataset to 85 genes and 493 transcript features. Gene-level omnibus tests were followed by stageR transcript-level adjustment at an overall FDR of 0.05. Pairwise dPSI was defined as the later-minus-earlier change in mean within-gene transcript proportion. Confirmed DTU required significant gene-level DRIMSeq results and at least one stageR-confirmed transcript.

Endpoint-specific DESeq2 tests quantify TSS or TES feature abundance relative to the corresponding endpoint-count layer; within-gene redistribution was evaluated separately by DTU analysis.

### Temporal trajectory classification

Gene trajectories were calculated as each gene’s fraction of evidence-filtered viral transcript counts and averaged across replicates. Peak classes reflected the time of maximum mean relative signal: 1 hpi, 2–4 hpi, 6–12 hpi, or 24 hpi. For profile classification, early abundance was the mean at 1, 2, and 4 hpi and late abundance the 24-hpi value. Genes with <25 total counts or no positive peak were low-signal; late-to-early ratios ≥1.5 and ≤1/1.5 defined late-rising and early-high profiles, respectively. Remaining genes peaking at 6 or 12 hpi were mid-peak; others were broad/stable. These are relative transcriptome profiles, not promoter-defined kinetic classes.

### Functional annotation and association analyses

OPGs were manually assigned to broad NCLDV-compatible functional classes using sequence homology, protein domains, reviewed annotations and literature. BLASTp v2.17.0+ was used against seven representative orthopoxvirus proteomes, InterProScan v5.77-108.0 for Pfam domains, RPS-BLAST v2.17.0+ for NCBI Conserved Domain Database matches, and DIAMOND blastp v2.2.0 against reviewed UniProtKB/Swiss-Prot proteins (49–53). Final assignments integrated these sources and relevant literature.

Copy-related OPGs, including inverted-terminal-repeat duplicates, were handled in a copy-aware manner. Associations of functional class with temporal profile, confirmed DTU, and high TSS/TES diversity were tested in feature-specific eligible gene sets. Temporal-profile analyses used the 139 main-ORF genes with assigned profile classes; DTU-related analyses were restricted to the DRIMSeq/stageR-testable main-ORF genes; and TSS/TES-diversity analyses used the corresponding endpoint-eligible gene sets. Pearson chi-square tests were used as exploratory omnibus screens across functional classes, and class-specific enrichment or depletion was assessed by one-versus-rest two-sided Fisher exact tests with Benjamini–Hochberg correction. The contribution of 5′ poly(A)-derived models was considered substantial at ≥100 total counts and ≥10% of gene-associated transcript signal. High TSS and TES diversity were defined as ≥4 and ≥6 retained features per gene, respectively. All class-level associations were treated as exploratory.

### Genomic-region and comparative conservation analyses

All 179 reference OPG features were retained for regional and comparative analyses. The conserved central region was defined as OPG048–OPG151; OPGs outside this interval were terminal. For functional summaries, genome-terminal localization was defined as within 25 kb of either genome end.

Comparative analyses included VACV Copenhagen (M35027.1), VACV Western Reserve (LT966077.1), cowpox virus Brighton Red (NC_003663.2), variola virus (NC_001611.1), ectromelia virus Moscow (NC_004105.1), camelpox virus M-96 (NC_003391.1), and taterapox virus Dahomey 1968 (NC_008291.1). Regional nucleotide effective similarity was alignment identity multiplied by aligned-region coverage; protein-level effective similarity was amino-acid identity multiplied by MPXV query coverage, summarized across comparator genomes by the median.

For the VACV-WR comparison, regional coding-feature conservation was summarized by mean effective OPG similarity, assigning 0 to missing homologous hits. Inferential protein-conservation analyses used median effective similarity across the seven comparators and excluded duplicated ITR features. Regional and functional differences were tested by Kruskal–Wallis tests. Functional- and temporal-class effects were additionally tested by Freedman–Lane-style residual permutation (54) on conservation ranks, stratified by genomic region; functional models adjusted for region and temporal models for region and functional class. Tests used 19,999 permutations, plus-one P values, and BH-adjusted pairwise comparisons.

### Cross-study comparison

For comparison with Picarone et al. (5), supplementary tables were imported and OPG identifiers harmonized with copy-aware mapping. Because that study used short-read gene-level measurements in a different cell model at 2, 24, and 48 hpi, whereas the present study used endpoint-filtered long-read-derived summaries from 1–24 hpi, timing agreement was assessed only at broad temporal resolution. The 37-gene comparison set was the deduplicated union of genes in Picarone et al.’s all-time top-20 table and additional genes explicitly highlighted in their main-text temporal examples; 35 were represented in the present analysis.

Published Picarone timing assignments were compared with present-study profile and peak-phase calls. Early Picarone genes were considered broadly concordant with present-study early-high or 1–4-h peak calls and different from present-study late-rising or 24-h peak calls; the inverse criteria were applied to Picarone intermediate/late genes. Other patterns were treated as partly comparable, and genes without a gene-specific Picarone assignment were not classified.

Yi et al. (6) were used qualitatively because available supplementary material lacked a viral temporal count matrix or gene-level differential-expression table suitable for harmonized reanalysis.

## Data Availability

CAGE-seq data are available through the European Nucleotide Archive under PRJEB60061. Direct cDNA and direct RNA sequencing data are available under PRJEB56841. Analysis scripts, processed tables, figure-building code, and package manifests are available at https://github.com/Balays/MPOX_PV424067.1. The 5,674-model discovery catalogue, transcript-level endpoint-support flags, and analysis-layer summaries are included with the supplementary package.

## Author contributions

This study is a reanalysis of previously published datasets; no new experiments were performed for the present work. G.Á.N. performed data curation, formal analysis and visualization and drafted the manuscript. B.K. carried out bioinformatics, data curation, formal analysis and interpretation of the data and contributed to drafting and revising the manuscript. Z.C. participated in the preparation of the manuscript. Á.D. participated in the preparation of the manuscript. I.P. participated in data analysis and manuscript preparation. D.R. contributed to methodology. J.H. participated in the preparation of the manuscript. G.K. contributed to conceptualization, resources and supervision. D.T. participated in study design, methodology and data analysis, and revised the manuscript. Z.B. conceived and designed the study, supervised and validated the work, participated in formal analysis and interpretation of the data, and contributed to drafting and revising the manuscript. All authors have read and approved the final manuscript.

## Ethics declaration

This study integrates previously generated transcriptomic data from earlier publications. No new sample collection, human or animal experimentation, or work with identifiable personal data was performed as part of the present analysis. Accordingly, additional ethical approval was not required for this work.

## Supplementary Figures Legends

Figure S1. Whole-genome overview of the transcript-region discovery catalogue. Of the 5,674 transcript models in the full catalogue, 2,392 informative models are displayed across four genome bands: 2,072 gene-assigned models reaching at least 2% mean within-gene usage at one or more sampled time point, and 320 unassigned or no-ORF models with nonzero dcDNA signal. Detailed coordinate-ordered regional views are provided in Supplementary File S1.

Figure S2. Candidate-model retention across analysis layers. Figure S2A, model counts for catalogue, gene assignment, main-ORF association, TSS/TES qualification, gene-level and transcript-usage sets. Figure S2B, the same values as percentages of the 5,674-model discovery catalogue.

Figure S3. Heatmap of the 30 highest-contributing endpoint-supported main-ORF genes. Values are row-scaled mean relative abundances across 1, 2, 4, 6, 12, and 24 hpi.

Figure S4. Highest-contributing endpoint-supported main-ORF genes. Ranked total counts for the 20 highest-contributing genes.

Figure S5. A, Temporal differential abundance detected by DESeq2 likelihood-ratio testing. Significant Main-ORF genes, TSS features and TES features at FDR < 0.05. B, Feature denominators for temporal differential-abundance testing. Tested Main-ORF genes, TSS features and TES features after count filtering.

Figure S6. Cross-study timing comparison with Picarone et al. Summary of 37 named Picarone genes, present-study temporal and DTU results. Categories are not mutually exclusive. Detailed gene-wise comparisons are provided in Table S5.

Figure S7. Relationship between supported TSS and TES feature diversity per gene. Each point represents a gene; selected loci with high supported endpoint multiplicity are labelled.

Figure S8. Time-resolved circular atlas of MPXV 5′-end architecture. Panels A–F show 1, 2, 4, 6, 12 and 24 hpi, respectively, with supported TSS clusters and log2-transformed dcDNA-seq coverage. Panel G shows pooled dRNA-seq as complementary native-RNA context.

Figure S9. Time-resolved circular atlas of MPXV 3′-end architecture. Panels A–F show 1, 2, 4, 6, 12, and 24 hpi, respectively, with supported TES clusters and log2-transformed dcDNA-seq coverage. Panel G shows pooled dRNA-seq as complementary native-RNA 3′-end and coverage context.

Figure S10. Temporal change in the relative abundance of candidate 5′ poly(A)-associated transcript models. Mean percentages of dcDNA-seq transcript counts assigned to candidate 5′ poly(A)-associated models and standard annotated models are shown across the six infection time points. Points represent means across three biological replicates; error bars indicate between-replicate variation.

Figure S11. Gene-level temporal dynamics of candidate 5′ poly(A)-associated transcripts. Heat map showing the mean proportion of these transcripts for each gene across the infection time course. Proportions were recalculated from per-sample counts of candidate 5′ poly(A)-associated and standard annotated transcript models and then averaged across biological replicates. Genes were required to have informative proportions in all three replicates at all six time points and at least 30 total classified transcripts at both 1 and 24 hpi. Rows are ordered by the change in the 5′ poly(A)-associated transcript proportion between 1 and 24 hpi, with the largest increases at the top. The complementary standard annotated transcript proportions are not shown because the two categories sum to 1.

Figure S12. Genes with the greatest candidate 5′ poly(A)-associated contribution. Genes are ranked using 5′ poly(A)-associated counts from the corrected catalogue and 18-library dcDNA time course; accompanying values summarize their contribution to the assigned gene signal.

Figure S13. Pairwise dPSI heatmap of stageR-confirmed transcript features. dPSI is later-time mean within-gene transcript proportion minus earlier-time mean; comparison labels therefore use later h − earlier h. The diverging scale reports the numerical dPSI range.

Figure S14. Comparative genomic organization and coding-feature similarity of MPXV and representative orthopoxviruses. Regional and OPG tracks show effective similarities rather than raw identity. Curated MPXV OPGs use the eight functional classes defined in Table S9.

## Supplementary Table Legends

Table S1. Complete 5,674-model discovery catalogue with origin, coordinates, gene/main-ORF assignments, endpoint-support flags and analysis-set eligibility.

Table S2. Main-ORF gene-level transcript-filter and count-matrix construction summary.

Table S3. Evidence-filtered main-ORF gene temporal summary for 139 loci.

Table S4. DESeq2 temporal differential-abundance summary for Main-ORF gene, TSS and TES layers.

Table S5. Published Picarone timing assignments compared with evidence-filtered present-study temporal and DTU results.

Table S6. Peak-refined CAGE-supported TSS feature summary with feature metadata and total adapter-correct dcDNA-seq counts.

Table S7. Origin-relevant LoRTIA dRNA-supported TES feature summary with feature metadata, LoRTIA_support_source, exact-support status and total adapter-correct dcDNA-seq counts.

Table S8. DRIMSeq/stageR differential-transcript-usage results. S8A contains gene-level results for the 85 fitted genes with test-specific p-value fields. S8B contains the 93 stageR-confirmed transcript features and their dPSI values.

Table S9. Evidence supporting the curated functional classification of MPXV OPGs, including homology, domain, reviewed-annotation, and literature information.

Table S10. OPGs with discordant broad functional classifications between the present study and Picarone et al.

Table S11. Integrated functional and transcript-architecture summary and functional-class association analyses. S11A, gene-level functional, temporal, architectural, genomic-position, and differential-transcript-usage summary for the 139 evidence-filtered main-ORF genes. S11B, functional-class association statistics, comprising exploratory omnibus Pearson chi-square tests and class-specific one-versus-rest two-sided Fisher exact tests performed in feature-specific eligible gene sets, with corresponding denominators, odds ratios, nominal P values, and Benjamini–Hochberg-adjusted P values.

## Supplementary File Legends

**File S1.** Regional transcript atlas of the discovery catalogue. The file contains 74 coordinate-ordered locus panels plus one unassigned/no-ORF panel. Of the 5,674 transcript models in the full catalogue, 2,392 informative models are displayed: 2,072 gene-assigned models reaching at least 2% mean within-gene usage at one or more sampled time-point, and 320 unassigned or no-ORF models with nonzero dcDNA signal. Together, these panels provide detailed regional views of the models summarized in Figure S1.

